# Single-cell RNA sequencing reveals that host Glutamine Metabolism Inhibition Enhances Macrophage Phagocytosis of *Mycobacterium tuberculosis*

**DOI:** 10.64898/2026.05.14.725218

**Authors:** Yingli Shi, Lesly Rodriguez Vicens, Micheal Teve, Barbara Slusher, William R Bishai, Sadiya Parveen-Nesson

## Abstract

Macrophages are crucial for host defense against the pathogen. However, pathogens such as *Mycobacterium tuberculosis (Mtb)* have evolved mechanisms to alter macrophage physiology and exploit these cells as their primary niche. *Mtb*-infected macrophages upregulate several metabolic pathways including glutamine metabolism. We previously showed that inhibiting glutamine metabolism with the pleiotropic glutamine metabolism antagonist prodrug JHU083 has dual antibacterial and immunomodulatory effects in a mouse model of tuberculosis. In the present study, using single-cell RNA sequencing and LS-MS/MS metabolomics, we showed that JHU083-mediated glutamine metabolism inhibition increased the population of interstitial macrophages in *Mtb*-infected lungs. JHU083 treatment also increased inflammatory signatures while lowering immunosuppressive markers on these macrophages. Metabolically, these macrophages exhibited marked depletion of complex lipids, accumulation of free fatty acids, and increased expression of transcripts associated with the β-oxidation pathway. Additionally, JHU083-treatment also improved phagocytic activity of macrophages, as measured by using fluorescent *E. coli* as a bait. In conclusion, JHU083-mediated glutamine metabolism inhibition metabolically reprograms macrophages, increasing both their lipid utilization as well as phagocytic activity, potentially driving their antimycobacterial activity that we had observed earlier.

## INTRODUCTION

Macrophages are professional phagocytes that serve as the first line of defense against invading pathogens ^1^. However, *Mycobacterium tuberculosis* (*Mtb*), one of the most successful intracellular bacterial pathogens, has evolved multiple strategies to survive within macrophages and evade their antimicrobial killing mechanisms ^2^. Following *Mtb* infection, macrophages undergo substantial metabolic reprogramming both in vitro and in vivo, and these metabolic alterations may promote intracellular bacillary survival and growth ^3^. However, the precise metabolic mechanisms underlying macrophage dysfunction during *Mtb* infection remain poorly understood.

*Mtb*-infected macrophages upregulate both glutamine uptake as well as expression of genes associated with glutamine metabolism in both mouse and human models of infection ^4-6^. In our previous study, we showed that blocking host glutamine metabolism with JHU083 significantly reduces bacillary burden in macrophages and mouse lungs ^7^. JHU083 is a pleiotropic glutamine metabolism inhibitor, a prodrug of glutamine antagonist, 6-diazo-5-oxo-L-norleucine (DON), which irreversibly inhibits multiple glutamine-utilizing enzymes, including glutaminases, glutamine synthetases, and several glutamine amidotransferases involved in the biosynthesis of purines, pyrimidines, coenzymes, hexosamines, and amino acids ^8,9^. We also showed that JHU083 administration potentiates effector T cell immunity and increases the frequency of interstitial macrophages in the lung ^7^. These macrophages also exhibited higher nitric oxide production than those treated with PBS or Isoniazid controls ^7^. Based on this data, we hypothesize that inhibition of glutamine metabolism increases macrophage antimycobacterial activity.

In the present study, using single-cell RNA sequencing-based *in silico* metabolomics coupled with traditional LC-MS/MS metabolomics analysis, we show that glutamine metabolism inhibition reprograms the inflammatory and metabolic state of macrophages, finally promoting their phagocytic activity.

## RESULTS

### 1. Single-cell RNA sequencing revealed 16 distinct immune populations in the Mtb-infected lungs

We reported earlier that JHU083-mediated host glutamine metabolism inhibition increases the frequency of interstitial macrophages (CD11b^+^ F4/80^+^) in the *Mtb*-infected lungs at 5-week post-infection, but not at 2-week post-infection, coinciding with the maximum bacterial clearance in the lungs ^7^. To dissect the phenotype of these late recruited interstitial macrophages, we decided to perform scRNA sequencing of *Mtb*-infected lungs with and without JHU083 treatment. Three groups of 129S2 mice were challenged with ∼75 CFU *Mtb* H37Rv, and then individual groups were treated with either 1 mg/kg JHU083 or 10 mg/kg rifampin (RIF) starting day 1 (**Fig 1A**). After 5 weeks of infection and treatment, mice were sacrificed from untreated (Un; n = 3), JHU083-treated (n = 3) and RIF-treated (n = 3) groups, lungs were harvested, single cell suspensions were prepared, and immune cells from each group were enriched using CD45^+^ magnetic bead-based enrichment method. Following enrichment, cells were loaded on to 10x Genomics Chromium Controller to capture mRNA on bar coded beads on approximately 10,000 individual cells per sample. cDNA libraries were prepared; quality checked and were sequenced at the depth of 50K reads per cell (**Fig S1**). The number of the cells captured per group ranged between 7111 to 8218 cells (**Table 1**). These CD45^+^ enriched cell populations primarily consisted of immune cells (Un = 93.96%, JHU083 = 98.27% and RIF = 86.67% of total cells). The top variable features were identified and used to perform UMAP-based cell clustering using the Seurat package in R ^10,11^. All non-immune cell clusters were removed *in silico* from the final analysis. Post UMAP clustering, we manually identified 16 prominent immune cells clusters of lymphoid (total 9 clusters) and myeloid lineages (total 6 clusters) (**Fig 2, Table 2**). The lymphoid lineage clusters included **(1)** CD4^+^ Naïve T cells (CD4_Na), **(2)** CD8^+^ Naïve T cells (CD8_Na), **(3)** Exhausted Effector CD4^+^ (CD4_Ex), **(4)** Terminally differentiated and exhausted effector CD4^+^ T cells (CD4_Tex), **(5)** Exhausted Effector CD8^+^ T cells (CD8_Ex), **(6)** Terminally differentiated and exhausted effector CD8^+^ T cells (CD8_Tex), **(7)** Regulatory CD4^+^ T cells (Tregs), **(8)** CD4^+^ CD8^+^ double-positive T cells (DPT) and, **(9)** mature B-cells (B). The myeloid clusters included (**9)** Proinflammatory M1-like macrophages (M1_Mac), **(10)** Pro-resolution M2-like macrophages (M2_Mac), (**11**) Classical monocytes (cMo), **(12)** Alveolar macrophages

**Fig 1.**
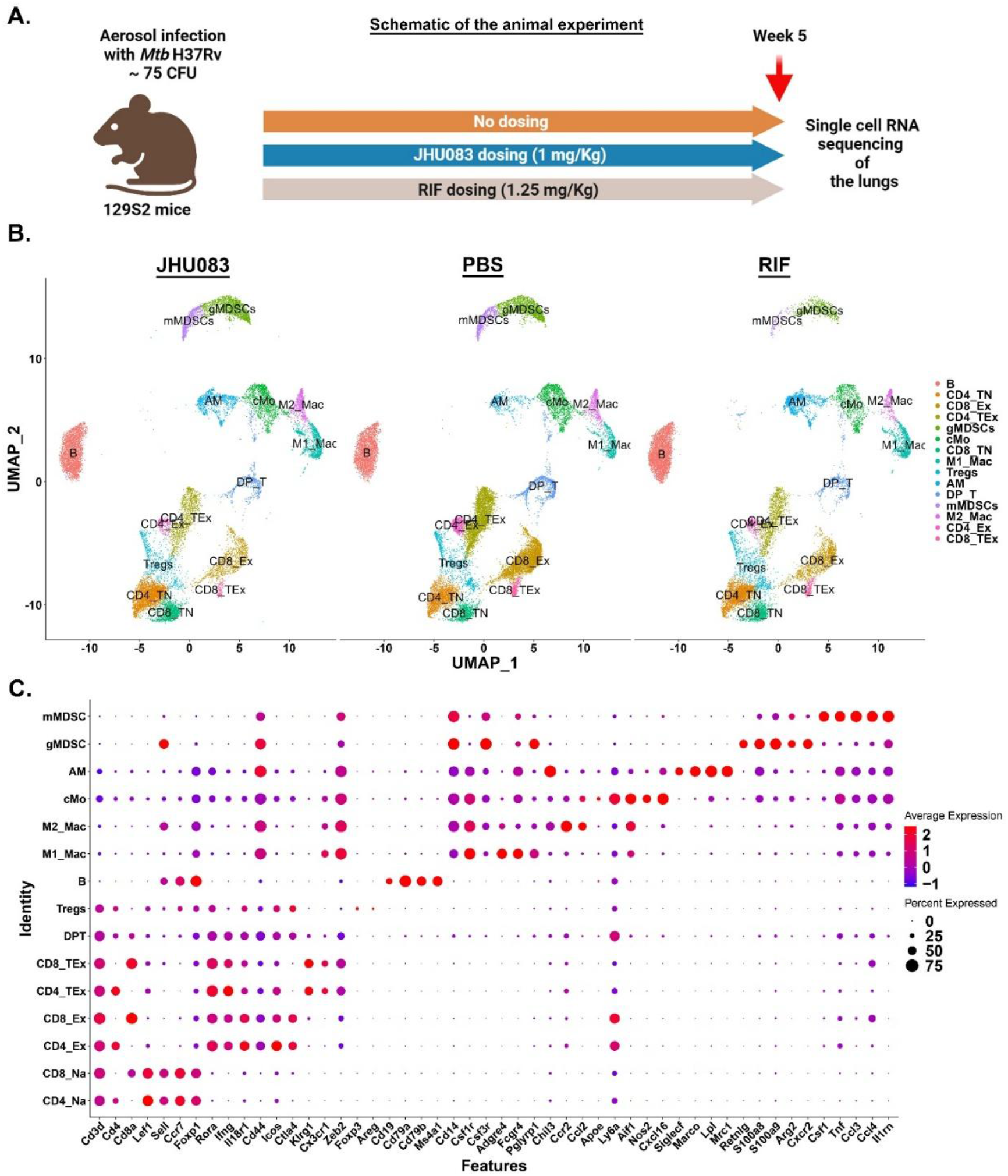
scRNA-seq transcriptional profiling of lungs. **(A)** Schematic representation of the murine experiment. 129S2 mice (n=10/group) were infected with ∼75 CFU of *Mtb* H37Rv. JHU083 or rifampin (RIF) was administered to the mice orally starting one day after treatment. 1 mg/kg was given daily for the first week, and then the drug was given on alternate days every week till the termination of the experiment. **(B)** UMAP plots showcasing all the immune cell clusters across in all three treatment groups. **(C)** Dot plot represents the key genes used to identify each specific immune cell cluster. The dot size represents the percentage of the cells expressing the specific transcripts in that cell cluster.

**Fig 2.**
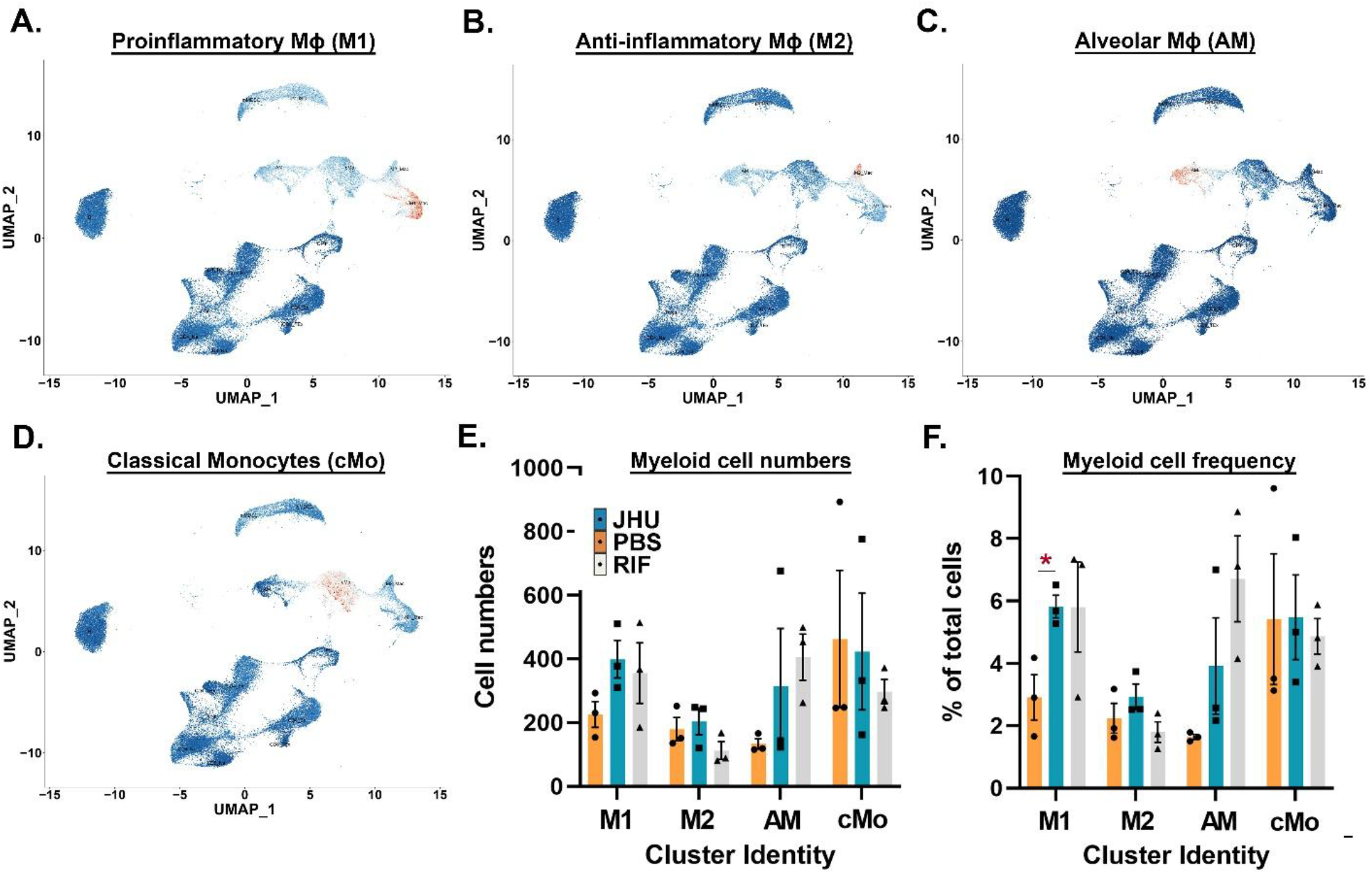
JHU083 administration enriches proinflammatory myeloid cells in the lungs. Feature plot showing the distribution of cluster-defining transcript expression on **(A)** proinflammatory macrophages (M1), **(B)** anti-inflammatory macrophages (M2), **(C)** alveolar macrophages (AM), and **(D)** classical monocytes (cMo). Bar graphs depicting **(E)** absolute number and **(F)** frequency of individual myeloid cell clusters in the dataset. Cell frequency was calculated as the percentage of total immune cells within in each dividual sample. The data is presented as Mean+SEM. Upaired t-test considering unequal distribution was to calculate statistical significance. * P <0.05.

(AM), **(13)** Granulocytic myeloid-derived suppressor cells (gMDSC) and **(14)** Monocytic myeloid-derived suppressor cells (mMDSC) (**Table 2**).

### 2. Host glutamine metabolism inhibition increased population of interstitial macrophages in the Mtb-infected lungs

For this study, we primarily focused on myeloid cell clusters and specifically interstitial macrophages (**Fig 2A-2D**). As shown in **Fig 2E and 2F**, JHU083-treated animals had significantly increased numbers and frequency of M1 macrophages in the lungs compared to untreated control mice lungs. While we found no difference in frequency of M2 macrophages, alveolar macrophages and monocyte clusters between JHU083-treated and PBS-treated mice, consistent with our earlier report. While RIF-treated animals exhibited no significant change in the frequency and number of M1 macrophages but exhibited a marked decline in the frequency of M2 macrophage cluster, most likely due to RIF’s differential mode of action compared to JHU083 (**Fig 2E and 2F**). Based on this data, we conclude that JHU083-mediated glutamine metabolism inhibition does increase interstitial macrophage population in *Mtb*-infected lungs.

### 3. JHU083 administration increases inflammatory signatures on M1 macrophages

We then performed DESeq2 based differential expression analysis of M1 macrophage cluster to investigate the effect of glutamine metabolism inhibition upon its transcriptional profile. We observed significant alterations in expression of genes associated with pathways crucial to macrophage physiology including inflammation, immunosuppression, phagocytosis and apoptosis (**Fig S2**). Using the AddModuleScore function in the Seurat package, we quantified the expression score of these pathways in individual macrophage clusters (see **Supplementary Table 1** for the list of transcripts). First, we observed a significant upregulation of inflammatory signatures on all macrophage clusters in JHU083-treatment groups compared to controls (**Fig 3A-D**). In contrast, M1-macrophage cluster from RIF-treated group exhibited substantially lower inflammatory signatures. At 5-weeks post-infection/treatment, both JHU083 and RIF-treated animals exhibit comparable levels of lung bacillary burden, the difference in the inflammatory signature between JHU083- and RIF-treated animals seems to be most likely driven by their differential mode of action (**Fig 3A-D**). As phagocytosis activity of macrophages is pivotal for the control of the *Mtb* proliferation, we then quantified the level of transcripts associated with phagocytosis and efferocytosis (**Supplementary Table 1**). There was substantial upregulation of the transcriptional signature associated with apoptotic cell clearance (**Fig 3E**), phagosome-lysosome fusion (**Fig 3F**) and overall phagocytosis (**Fig S3**) in JHU083-treated groups compared to the untreated controls. M1-macrophage cluster from JHU083-treated groups also exhibited modest but significant decrease in the expression score of immunosuppression (**Fig 3G**). Taken together, these observations suggest that JHU083-mediated glutamine metabolism inhibition reprograms macrophage cluster towards a proinflammatory and phagocytic phenotype with a lowered immunosuppressive activity in the hyperacute model of *Mtb* infection.

**Fig 3.**
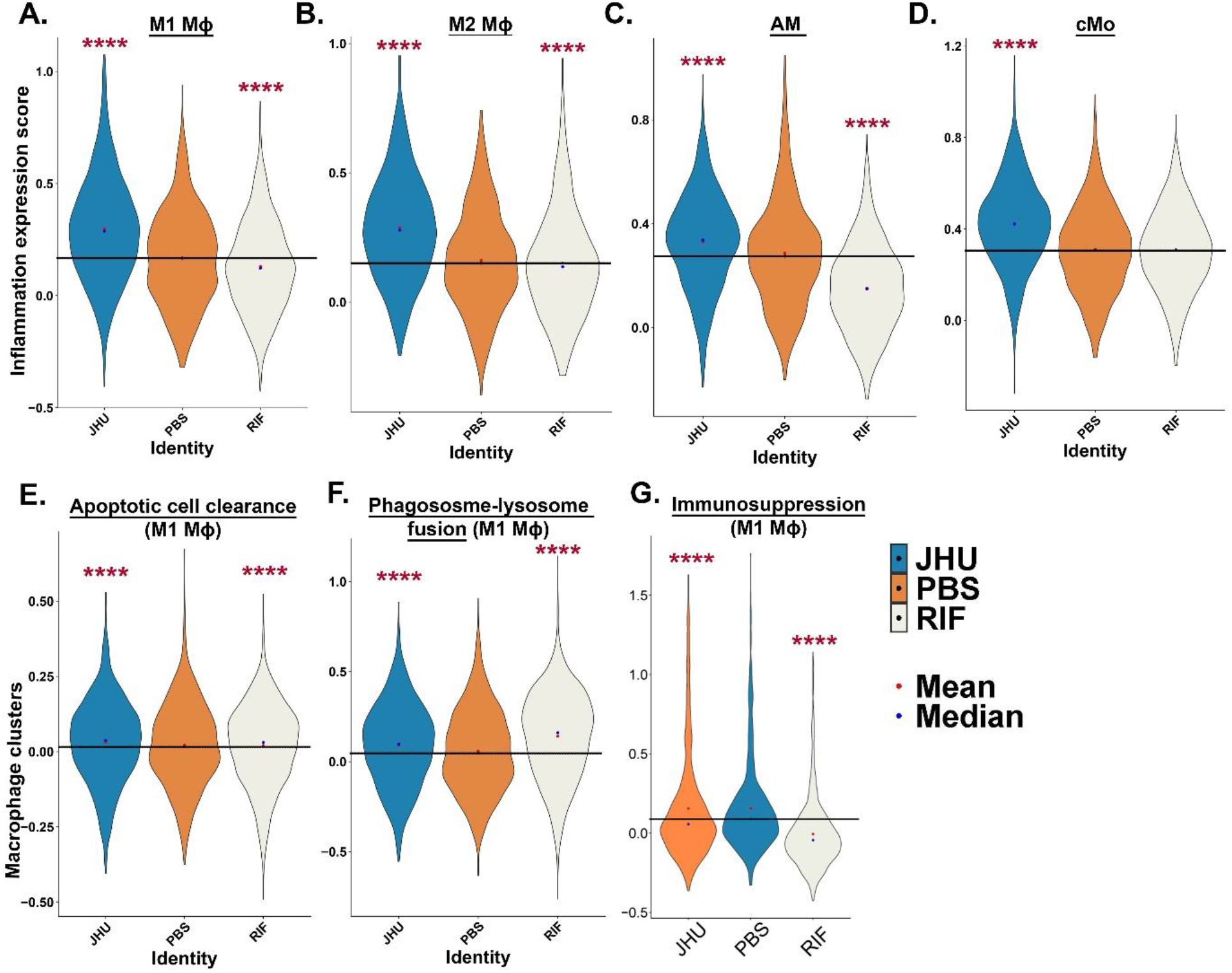
JHU083 treatment alters expression signatures of inflammation, phagocytosis and immunosuppression. Violin plot depicting expression score of inflammation upon **(A)** proinflammatory macrophages, **(B)** anti-inflammatory macrophages, **(C)** alveolar macrophages, and **(D)** classical monocytes. We then calculated the expression score of two processes associated with phagocytosis: **(E)** apoptotic cell clearance, **(F)** phagosome-lysosome fusion and, **(G)** immunosuppression on the macrophage clusters alone. The expression score was calculated using the AddModuleScore module in the R package. The transcripts used for the expression score calculations are listed in **supplementary table 2**. The solid line represents the value corresponding to the mean value of the untreated control samples (PBS). Mean (red dot) and median (blue) values were plotted using ggplot2. ANOVA was used to calculate statistical significance. **** P<0.0001.

### 4. In silico metabolomics analysis identifies a unique myeloid cell-specific metabolic reprogramming in JHU083-treated group

Since we reported earlier that JHU083 reprograms several metabolic pathways in the *Mtb* infected lungs and tumor microenvironment (Parveen et al., Leone et al., oh et al.), we further investigated the effect of JHU083 administration upon cellular metabolome of M1-macrophages. Using the AddModuleScore function in the Seurat package, we quantified the expression score of various metabolic pathways (see **Supplementary Table 2** for transcripts) after binning the data into two distinct subsets; (1) total CD45^+^ clusters and, (2) all myeloid clusters. As shown in **Fig 4A**, we focused on various metabolic pathways feeding into cellular ATP production. Both subsets exhibited significant downregulation of glycolysis and pentose-phosphate pathways in JHU083- and RIF-treated groups, compared to untreated control. As both glycolysis and PPP provide intermediates for the tricarboxylic acid (TCA) cycle, ultimately feeding into oxidative phosphorylation generating ATP, we quantified the TCA cycle and oxidative phosphorylation expression score. TCA cycle was significantly upregulated in myeloid clusters in both treatment groups compared to the control. In comparison, oxidative phosphorylation remained upregulated in both subsets as well as treatment groups compared to the control (**Fig 4B and 4C)**. Finally, we explored fatty acid oxidation (β-oxidation) and amino acid utilization as alternative sources of acetyl CoA for the TCA cycle. We detected higher β-oxidation scores in the myeloid clusters in both JHU083- and RIF-treated mice compared to control mice (**Fig 4B and 4C**). Overall, *in silico* metabolomics analysis suggests that in presence of JHU083-mediated glutamine metabolism inhibition, myeloid cells such as macrophages seem to rely on lipids as their preferential energy source. And this complex metabolic rearrangement in the lung immune microenvironment, potentially leads to an enhanced antibacterial activity of macrophages.

**Fig 4.**
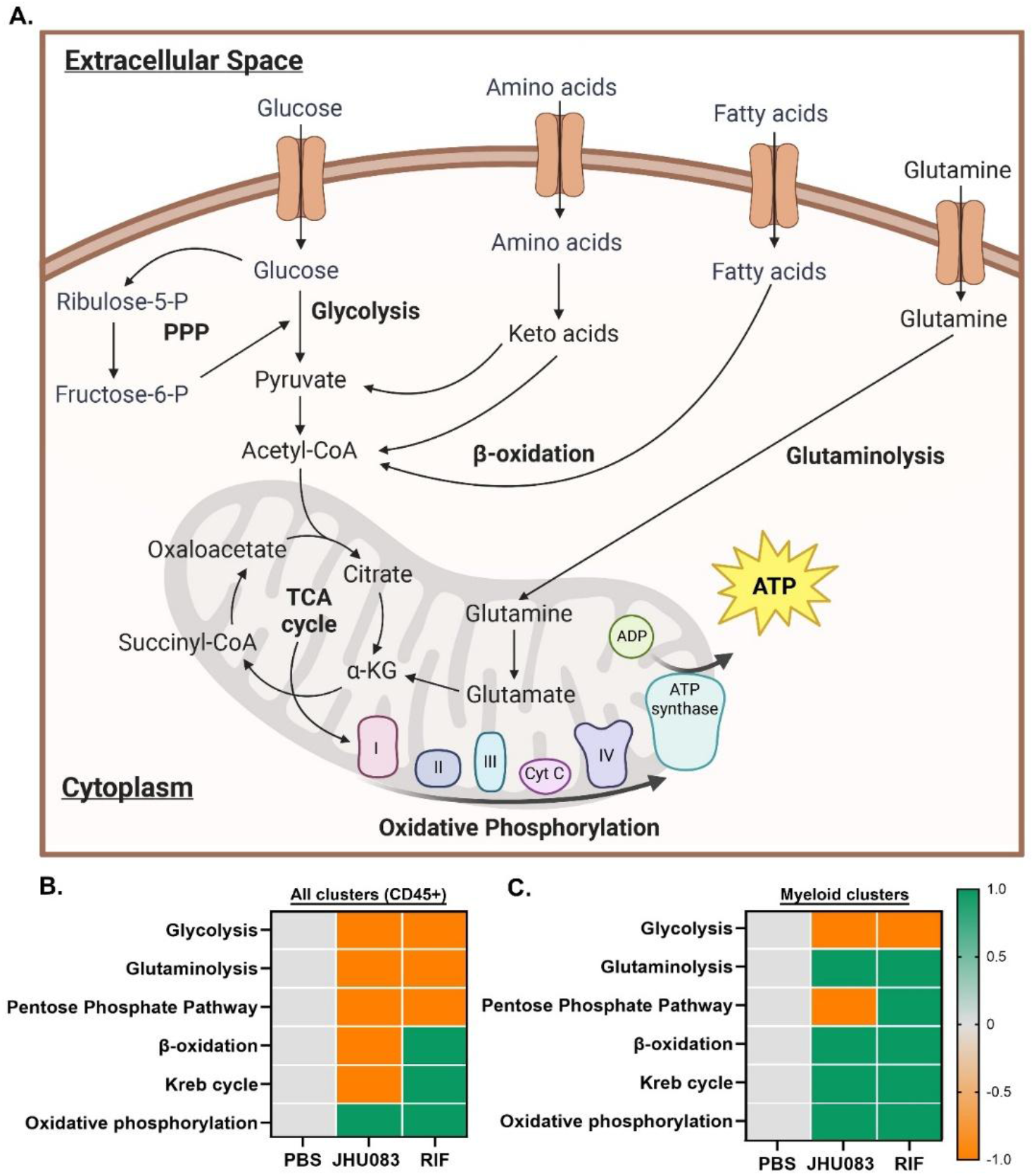
JHU083 administration leads to immunometabolic reprogramming in myeloid cells. **(A)** Schematic highlighting the major energy-generating pathways present in the cell. The pathways analyzed in the study are highlighted in bold letters. Heatmap representing the relative expression score of various metabolic pathways in JHU083- and RIF-treated groups compared to control groups (PBS) in **(B)** all CD45^+^ cells and **(C)** only myeloid cells (consisting of AM, M1, M2 and cMo clusters). The expression score was calculated using the AddModuleScore module in the R package. The transcripts used for the expression score calculations are listed in **supplementary table 3**. All CD45^+^, and myeloid clusters were subsetted into two datasets for expression score calculations. Zero value was assigned to the control group for all pathways; the expression score of 0, +1, and -1 was assigned to equal, higher, or lower values than the control groups. Only the changes with P<0.05 are plotted here.

### 5. LC/MS based metabolomics confirmed depletion of lipids and amino acids in JHU083-treated M1 macrophages

To confirm the substantial metabolic remodeling observed in myeloid cells using *in silico* metabolomics, we then investigated the effect of JHU083 administration upon cellular metabolome of primary macrophages. To do so, isolated BMDMs from 129S2 mice, infected with *Mtb* H37Rv and treated them with PBS, JHU083 and Isoniazid (INH) for 24 and 48 hours. We found significant differences in the level of metabolites extracted from macrophages treated with JHU083 for 24 hours compared to PBS-treated controls. There was modulation in the level of 156 metabolites in JHU083-treated macrophages compared to the untreated controls (P < 0.05), 24-hour post-treatment. Minimal changes in the metabolite levels at 48-hour time-point were noted (data not shown). Most importantly, we observed a 26% decrease in the intracellular level of glutamine consistent with JHU083-mediated glutamine metabolism inhibition (**Fig 5A**). We noted no change in the level of glutamate in these macrophages (**Fig 5B)**. In addition, we also noted significant decline in several other amino acids such as asparagine, proline, arginine, and alanine, and amino-acid related metabolites such as ornithine, acetyl-aspartate, taurine, glutaminylmethionine (**Fig S4**) possibly indicating either higher utilization of amino acids or their declining import. This is consistent with our earlier report wherein we observed depletion of several amino acids in the lungs of *Mtb*-infected mice treated with JHU083 ^7^.

**Fig 5.**
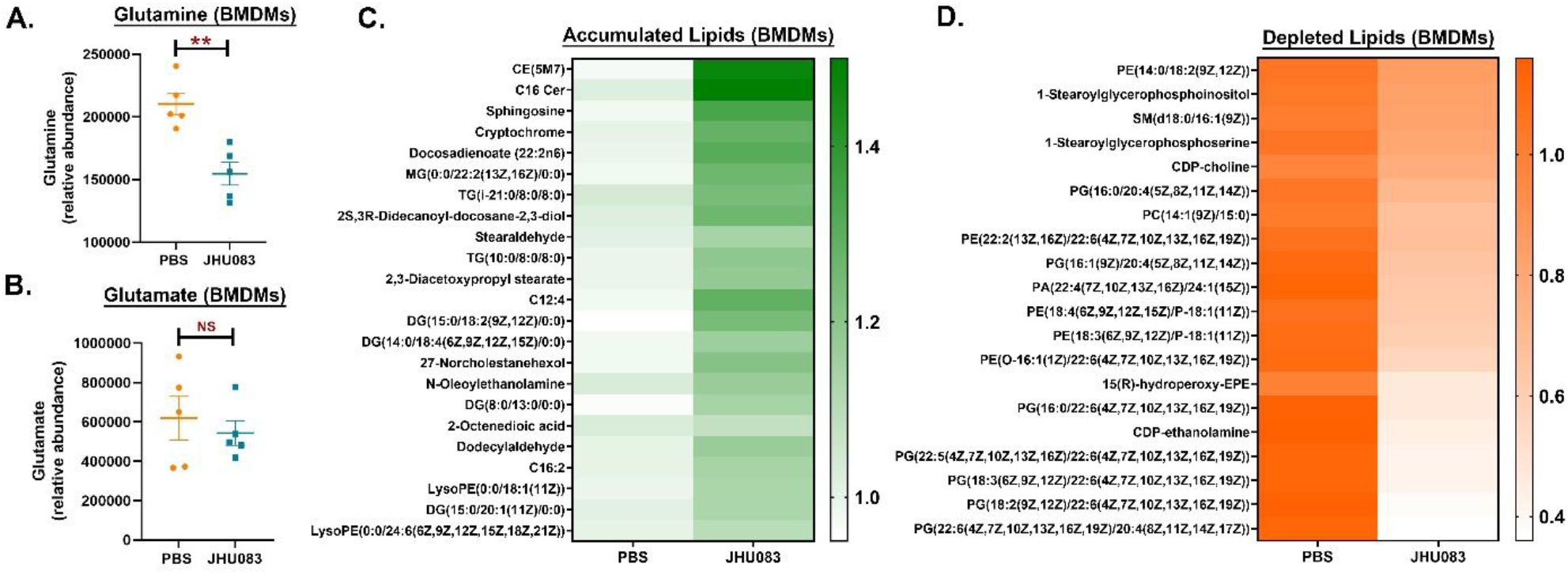
LC-MS/MS metabolomics revealed metabolic reprogramming in *Mtb*-infected macrophages upon JHU083 treatment. As described in “Methods”, *Mtb*-infected BMDMs were treated with JHU083, or PBS for 24 hours, and metabolites were extracted from macrophages using methanol as described in the “Methods.” Metabolite levels were normalized to cell number and then to untreated controls. We detected **(A)** significant changes in glutamine but no change in glutamate level in macrophages. Heatmap representing the top lipid metabolites that were either **(C)** enriched or **(D)** depleted in the macrophages treated with JHU083 (right-lane) compared to untreated group (left lane). Metabolite abundance was normalized to the number of macrophages used for extraction and to untreated controls. Statistical significance was determined using a two-tailed Student’s t-test assuming unequal variance. Data are representative of two independent experiments. ** P<0.01.

However, the most striking phenotype which we observed was the accumulation of free fatty acids concomitant with depletion of complex lipids such as phosphatidylglycerols in the JHU083-treated infected macrophages (**Fig 5C and 5D)**. The data is consistent with upregulation of β-oxidation expression scores in myeloid cells as seen with *in silico* metabolomics (**Fig 4C)** and indicates that JHU083 administration infected macrophages might preferentially rely on complex lipids and amino acids as the fuel source to generate sufficient energy when they could not access glutamine metabolism and glycolysis.

### 6. JHU083 administration increases phagocytosis activity of M1 macrophages

So far, our data suggested that JHU083-treatement induces distinct metabolic reprogramming of M1 macrophages and increases their proinflammatory phenotype. We have already reported earlier that JHU083 treatment of M1 macrophages reduces bacillary burden by 0.7 log in ex vivo macrophage infection model ^7^. To decipher how metabolic reprograming is leading to improved bacterial clearance by M1 macrophages, we evaluated effect of JHU083 administration upon the phagocytic activity of M1-like macrophages, using fluorescent *Escherichia coli* (*E. coli*) cells as bait. We observed a significant difference in the phagocytosis activity in JHU083-treated BMDMs compared to PBS-treated controls (**Fig 6A**). To confirm that this observation is not specific to the strain of mice, we tested the effect of JHU083 on the phagocytic activity of BMDMs isolated from C57BL/6J mice as well and found that JHU083-mediated glutamine metabolism inhibition improves phagocytic activity of macrophages independent of the strain background (**Fig 6B**). Based on this data, we conclude that host glutamine metabolism inhibition improves phagocytic activity of macrophages, potentially driving the antimycobacterial activity of macrophages described earlier.

**Fig 6.**
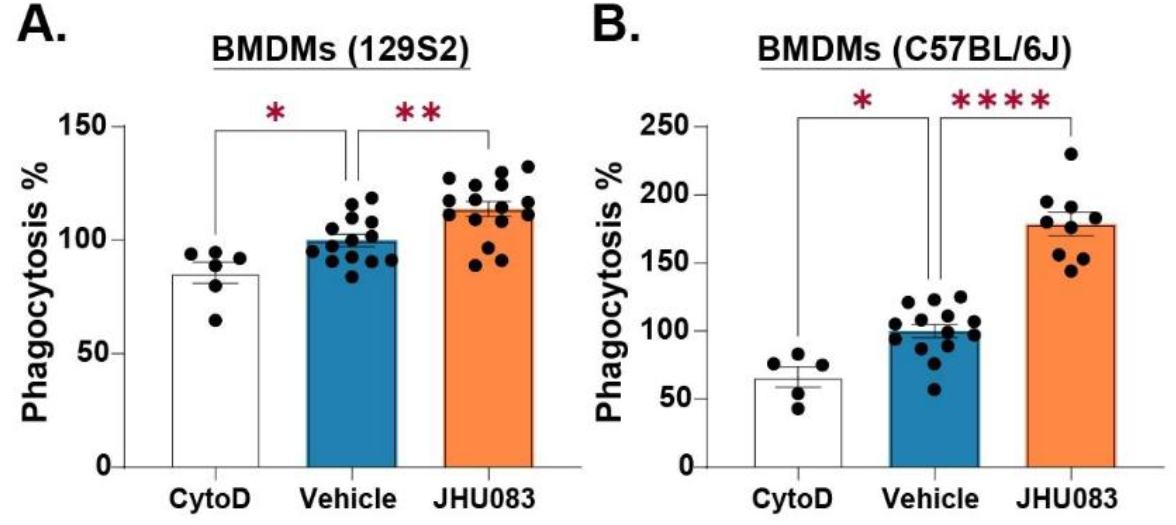
JHU083 treatment promotes phagocytic activity of macrophages. IFNγ-activated BMDMs from 129S2 or C57BL/6J mice were treated with JHU083 (10 μg/mL) or vehicle control prior to incubation with heat-inactivated green *E. coli*. Phagocytosis was quantified after 4 hours by fluorescence measurement following quenching of extracellular signal. Cytochalasin D (20 μM) was used as a negative control. Data are shown as mean ± SEM. Statistical significance was calculated using a two-tailed Student’s t-test. * P<0.05. ** P<0.01. **** P<0.0001.

## DISCUSSION

In our present study, we show that glutamine metabolism inhibition increases the population of interstitial macrophages in the *Mtb*-infected lungs. These macrophages exhibit increased proinflammatory and phagocytic behavior. Metabolomics analysis suggests that these macrophages utilize complex lipids as energy sources; as evidenced by depletion of complex lipids, accumulation of free fatty acids and increased expression of transcripts associated with β-oxidation. Overall, our study identifies a critical metabolic shift initiated by glutamine metabolism inhibition in macrophages which then leads to their enhanced phagocytic activity. Interestingly, while our observations help explain the therapeutic benefit exerted by JHU083, they also highlight an important paradox in the field. On one hand, glutamine plays an essential role in macrophage growth, proliferation, differentiation, and physiological function ^5,12,13^. On the other hand, pathogens such as *Mtb* upregulate glutamine metabolism in macrophages ^4,6^, potentially impairing macrophage antibacterial activity and promoting establishment of infection. In the present study, we observed decreased glutamine concentrations in macrophages, consistent with glutamine metabolism inhibition; however, transcriptionally, we observed increased abundance of transcripts associated with glutamine metabolism. Glutamine has also been shown to be critical for the inflammation-resolving capacity of M2 macrophages ^14^, and it will be interesting to examine the effects of JHU083 on M2 polarization in the context of *Mtb* infection. While glutamine metabolism targeting has been shown promise in several preclinical models of cancer^9,15,16^, host glutamine metabolism targeting as a host-directed therapy for TB remains in its infancy. We were the first group to demonstrate that inhibition of host glutamine metabolism exerts both antibacterial and immunomodulatory effects during *Mtb* infection ^7^. In our previous study, the therapeutic efficacy of JHU083 in murine TB appeared to be primarily host-mediated, as (1) JHU083 failed to confer protection in immunocompromised mice and (2) lung concentrations of JHU083 remained approximately 4,000-fold below the in vitro MIC against *Mtb* ^7^. We also observed selective enrichment of interstitial macrophages in the lungs following 5 weeks of infection and treatment ^7^; however, their phenotype could not be comprehensively characterized by flow cytometry alone.

In the present study, we utilized scRNA sequencing, *in silico* metabolomics, and LC-MS/MS-based metabolomics to define the effects of JHU083 on lung interstitial macrophages both in vivo and in an ex vivo macrophage model. An earlier study has demonstrated that glutamine is crucial for M1 polarization of *Mtb*-infected macrophages. They also showed that chemical or genetic inhibition of GLS1, a key enzyme that converts glutamine to glutamate, promoted *Mtb* growth in macrophages^5^. Qi et al. demonstrated that DON reshapes macrophage responses in a context-dependent manner. In M1 macrophages, DON did not uniformly enhance activation: during M1 differentiation, it increased inflammatory cytokine expression and phagocytic activity despite reducing MHC-II and CD68, whereas in fully polarized M1 cells it increased MHC-II/CD68 with only modest cytokine induction and no further improvement in phagocytosis. Notably, the DON-induced increase in MHC-II in fully polarized M1 macrophages required extracellular glutamine, as this effect was largely lost under glutamine-depleted conditions. By comparison, DON more consistently suppressed M2/TAM-like macrophages by reducing immunosuppressive features and increasing their susceptibility to viability loss and apoptosis. Together, these findings suggest that DON may shift macrophage balance primarily by weakening M2 programs while preserving selected M1-associated functions ^17^. The discrepancy among these studies could be due to the pleiotropic nature of JHU083 as a glutamine metabolism inhibitor prodrug as JHU083 irreversibly blocks several glutamine utilizing enzymes beyond GLS1^9^. Future studies examining glutamine uptake and pathway-specific metabolic flux may help resolve these discrepancies.

As JHU083 is already being evaluated in clinical trials for advanced-stage fibrolamellar carcinoma (NCT07430202, NCT06027086) and non-small cell lung cancer (NCT07249372), either as a monotherapy or in combination therapy, its clinical translation potential is substantial. However, before advancing toward clinical application in tuberculosis, the therapeutic efficacy of JHU083 must be evaluated during both the acute and chronic phases of TB. Additionally, given the primarily pulmonary presentation of TB, intranasal delivery of JHU083 may represent an attractive strategy to enhance drug delivery to the lungs while minimizing potential off-target effects, particularly considering the essential role of glutamine metabolism in normal human physiology. Overall, our study identifies host glutamine metabolism as an attractive target for host-directed therapy against TB.

### Limitation of the study

For metabolic studies, metabolic pulse-chase experiments will be necessary to accurately assess metabolic flux, as depletion of a metabolite may result from either increased utilization or reduced uptake. However, in the case of β-oxidation, we observed not only depletion of complex lipids, but also accumulation of free fatty acids along with increased expression of β-oxidation-associated transcripts, increasing our confidence in the finding that JHU083 treatment increases β-oxidation in macrophages infected with *Mtb*. Additional pulse chase experiments are being pursued by these authors to validate the findings further.

## MATERIALS AND METHODS

### Animal Infection Study

Female 129S2 mice (6–10 weeks old; n =10/group) were obtained from Charles River Laboratories. Mice were infected with ∼200 CFU of *Mycobacterium tuberculosis* H37Rv via aerosol infection. Sequencing-verified frozen stocks of *Mtb* H37Rv were used for all infection studies. One day post-infection, mice were randomly assigned to one of three treatment groups: (1) PBS control, (2) JHU083, or (3) rifampin (RIF). JHU083 was administered daily dosing consisting of 1 mg/kg/day during the first week followed by 0.3 mg/kg/day thereafter. Rifampin was administered orally at 12.5 mg/kg/day as a positive control. Only female mice were used to facilitate co-housing and minimize the number of animals required per experimental group.

### Single cell suspension of lung using enzymatic dissociation

Lungs were collected in 5 mL MACS Tissue Storage Solution (Cat: 130-100-008; Miltenyi Biotec) and stored at 4 °C until processing. The lungs were dissected into individual lobes, and single-cell suspensions were prepared using the mouse lung dissociation kit (Cat: 130-095-927; Miltenyi Biotec) and GentleMACS™ Dissociator (Miltenyi Biotec) according to the manufacturer’s instructions. Cell suspensions were incubated with ACK lysis buffer (Cat: A1049201; Gibco) at room temperature for 2-5 min to lyse red blood cells, followed by washing with RPMI complete media and resuspension in the appropriate volume of the same media. Cell viability was assessed using trypan blue staining.

### ScRNA sequencing and analysis

For performing scRNA sequencing, Chromium Next GEM Single Cell 3’ GEM, Library & Gel Bead Kit v3.1, 10X Genomics was used. Lung single cell suspension was enriched for CD45^+^ cells using EasySep™ Mouse CD45 Positive Selection Kit (Cat: 18945; Stem Cell Technologies). The final suspension consisted of ∼90% CD45^+^ cells of more than 90% viability. For capturing a total of 10,000 cells per sample, 16,500 cells per well per sample were loaded on a 10x Chromium controller. cDNA and library construction was performed following the manufacturer’s protocols.

mRNA libraries were sequenced at 50,000 reads per sample depth on an Illumina Novaseq by Experimental and Computational Genomics Core at Johns Hopkins University. ScRNA seq reads we processed with Cell Ranger v3.1 software to quantify transcript counts per cell. Quantification was performed using the STAR aligner against the *Mus musculus* transcriptome. For downstream processing and visualization, Seurat V3 package in RStudio was used. Cells with less than 500 features or more than 5% mitochondrial DNA were omitted from further analysis. We used top 30 PCAs at 0.6 resolution to generate cell clusters. We omitted clusters (1) not expressing CD45 (*Ptprc*) or (2) less than 2% frequency of the total cells (with two exceptions of alveolar macrophage and terminally exhausted CD8^+^ T cell clusters). Up to top 50 genes were used to identify the clusters manually. The clusters were also compared with each other to fine-tune the cluster nomenclature. Post naming the clusters, the final Seurat object was merged and DEseq2 package ^18^ was used to identify differentially expressed genes among the treatment groups. The differentially expressed genes were plotted using EnhancedVolcano package^19^ (P<0.05; average log fold change > 0.25). Top 30-60 upregulated and downregulated genes (P<0.05; average log fold change > 0.25) were used to analyze the differences among the treatment groups. The expression signature was calculated using AddModuleScore package ^20^ in R.

### BMDM isolation

BMDMs were generated from 8-12-week-old female C57BL/6J (Jackson Laboratoy) or 129S2 mice (Charles River Laboratories) mice as previously described ^21^. Bone marrow cells were isolated from femurs and cultured for 6 d in RPMI 1640 medium supplemented with GlutaMAX (1×) and 10% L929-conditioned medium as a source of M-CSF. After differentiation, BMDMs were plated in 12-well plates at 0.25 × 10^6^ cells/well for metabolite extraction, or in 96-well plates at 0.05 × 10^6^ cells/well for phagocytosis assays. Cells were pretreated with recombinant IFN-γ (20 ng/ml) for 24 h before drug treatment.

### BMDM infection and Metabolite extraction

Infection studies were performed as we described earlier ^7,21^. Briefly, isolated BMDMs from 129S2 mice were polarized with 20 ng/mL IFNγ for 24 h and infected with *Mtb* H37Rv at an MOI of 2 for 4 h. Cells were washed and incubated with 200 μg/mL gentamicin to eliminate extracellular bacteria, followed by treatment with JHU083 (10× MIC), or INH (32× MIC). After 24-and 48-hours post-infection and drug treatment, BMDMs were washed three times with freshly prepared 75 mM ammonium carbonate buffer (pH 7.4) and metabolites were extracted using cold (-20 °C) acetonitrile:methanol:water (40:40:20, v/v/v). Following a 10-min incubation at -20 °C, the extraction was repeated, and cell lysates were collected by scraping and pooling. Extracts were centrifuged at 14,000 rpm for 10 min at 4 °C, and the resulting supernatants were collected, filtered via 0.22 um acetate filter and stored at -20 °C until analysis. Parallel wells were processed for cell counting to normalize metabolite abundance across samples.

### LC MS/MS Analysis of metabolites

LC MS/MS analysis was performed as described earlier ^7^. Briefly, Dried metabolite extracts were resuspended in 50% acetonitrile and analyzed using an LC-MS platform consisting of an Agilent 1290 Infinity II UHPLC coupled to a Bruker timsTOF Pro II mass spectrometer equipped with an IonBooster ESI source. Metabolites were separated on a Waters XBridge BEH Amide HILIC column (2.1 × 150 mm, 1.7 μm) using a nonlinear gradient of water + 0.1% formic acid (solvent A) and acetonitrile + 0.1% formic acid (solvent B) from 99% to 45% B over 25 min at a flow rate of 0.2 mL/min. MS data were acquired in negative ion mode with auto MS/MS over a mass range of 70–1100 m/z. Data acquisition and analysis were performed using Compass HyStar 5.1 and TASQ 2022 software, respectively. Metabolites were identified using an in-house database with HILIC-based retention times. Due to structural similarities among certain metabolites, some analytes could not be definitively distinguished.

### Phagocytosis assay

Macrophage phagocytosis was assessed using a green *E. coli*-based phagocytosis assay kit (Cat: ab235900; Abcam) according to the manufacturer’s instructions. BMDMs were isolated from either 129S2 or C57BL/6J mice, IFNγ-activated and then treated with JHU083 (10 μg/ml) or vehicle control for 1 h before the addition of heat-inactivated green *E. coli*. After 4 h of incubation with green *E. coli*, cells were washed twice to remove noninternalized bacteria, and extracellular fluorescence was quenched according to the manufacturer’s protocol. Cytochalasin D (20 μM, Cat: C8273-1MG; Sigma-Aldrich) was used as a negative control. Fluorescence was measured at excitation/emission (Ex/Em) 490/520 nm using a SpectraMax iD3 multi-mode microplate reader (Molecular Devices).

## STATISTICS AND REPRODUCIBILITY

No statistical methods were used to predetermine sample size. All experiments were randomized, and all collected data are reported in the manuscript. No data were excluded from analysis. Following aerosol infection, animals were randomly assigned to experimental groups. Investigators were not blinded to group allocation during experiments or outcome assessment. Data was analyzed using GraphPad Prism 11. Comparisons between two groups were performed using unpaired two-tailed Student’s t tests. A P value < 0.05 was considered statistically significant.

## DATA SHARING

All scRNA-seq datasets, code used for scRNA-seq analysis, metabolomics datasets etc. will be made publicly available following the acceptance of this preprint in a peer-reviewed journal. Please reach out to the SPN for further information.

### ACKNOWLEDGEMENT

SPN gratefully acknowledge the support of TRAC developmental award (JHU TRAC NIH/NIAID Fund P30AI18436), the CoBRE Center for Biomedical and Laboratory Development Research Project Leader Award (NIH/NIGMS 5P20GM130555), the CoBRE Center for Applied Immunology and Immunotherapy Startup funds and Pilot Award (NIH/NIGMS 5P20GM134974). WRB acknowledges NIH/NIAID R01AI155602 funds. We acknowledge Experimental and Computational Genomics Core at Johns Hopkins for providing sequencing, genome annotation and Cell ranger pipeline services for ScRNAseq analysis. We acknowledge General Metabolics (Boston, MA) for providing metabolomics services. YS is a mentee of the NIH/NIGMS Career MODE Program (R25GM143298).

## AUTHOR CONTRIBUTIONS

WRB and SPN conceptualized the study and designed the research approach; BSS provided JHU083 and critical input for the study; YS, LRV, MK and SPN performed experiments; SPN performed scRNA seq studies and data analysis; YS, WRB and SPN analyzed data; YS, WRB and SPN wrote the manuscript; YS, LRV, MT, BSS, WRB and SPN critically reviewed the manuscript.

## COMPETING INTERESTS

YS, LRV, MT, WRB and SPN declare no conflict of interest. BSS is an inventor on multiple Johns Hopkins University patents related to glutamine antagonist prodrugs, including JHU083, and their therapeutic applications. These technologies have been licensed to Dracen Pharmaceuticals. R B.S.S. is co-founders of and hold equity in Dracen Pharmaceuticals. These arrangements have been reviewed and approved by Johns Hopkins University in accordance with institutional conflict-of-interest policies. The authors declare no additional competing interests.

---

**Fig S1.**
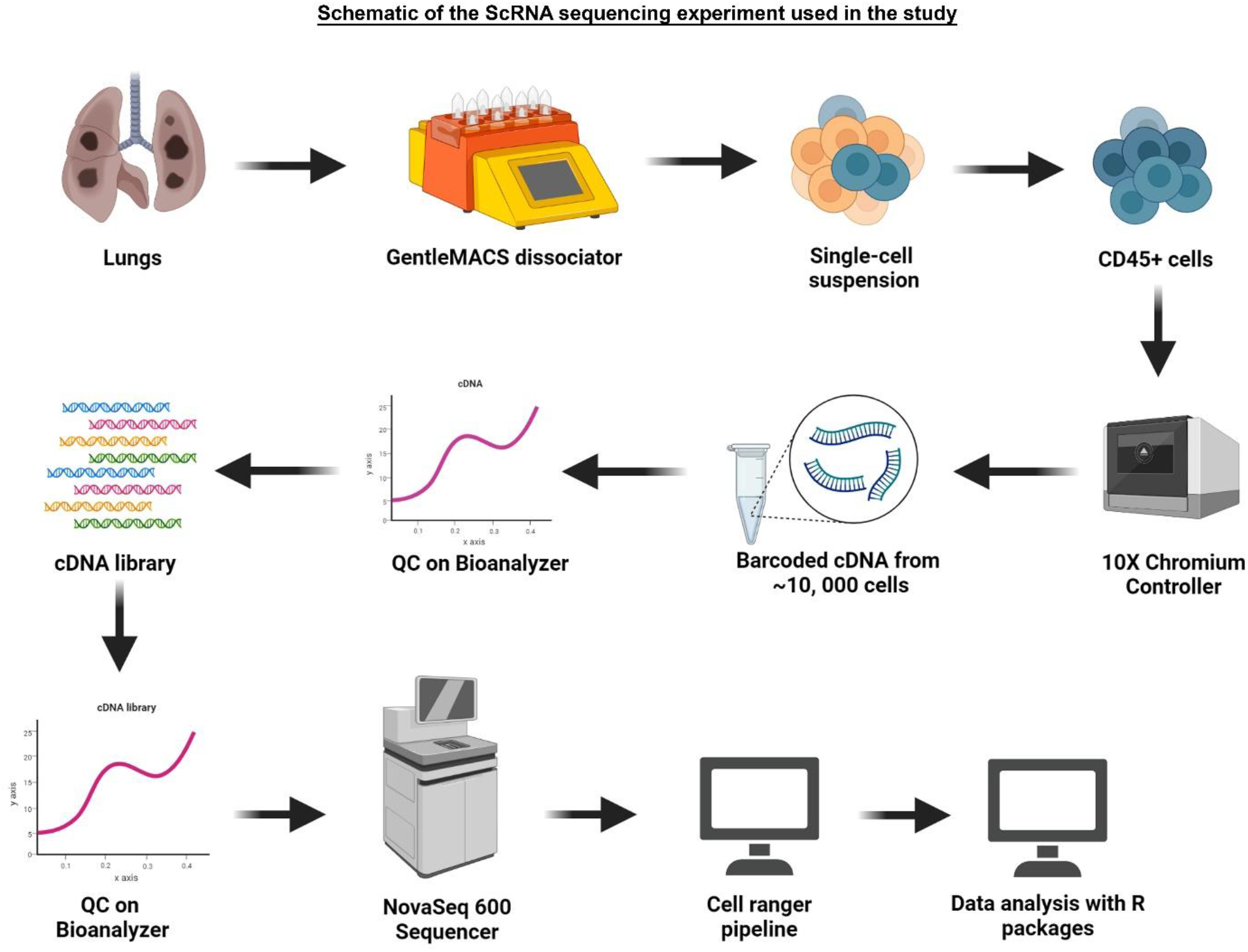
Experimental workflow used for scRNA sequencing experiment.

**Fig S2.**
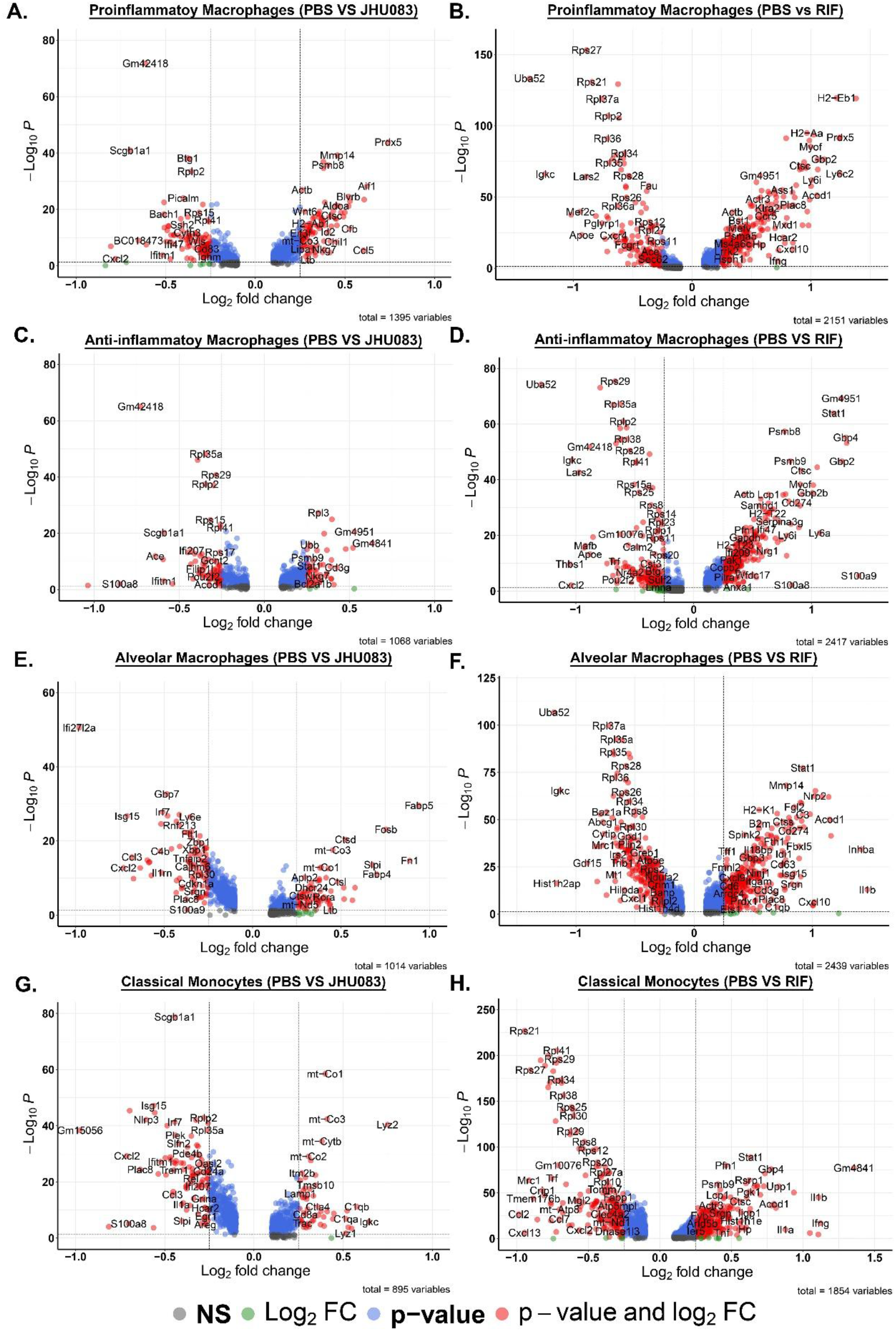
Scatter plot representing differentially expressed transcripts in myeloid cell clusters from both JHU083- and RIF-treated groups compared to the untreated control. The differential expression analysis was performed using the DESeq2 package in R of **(A)** proinflammatory macrophages, **(C)** anti-inflammatory macrophages, **(E)** alveolar macrophages, and **(G)** classical monocytes in JHU083-treated groups compared to the untreated controls (PBS). **(B)** Proinflammatory macrophages, **(D)** Proinflammatory macrophages, **(F)** alveolar macrophages, and **(H)** classical monocytes in RIF-treated groups compared to the untreated controls (PBS) are also plotted here. A P-value of 0.05 and an average log fold change of 0.25 were used as the analysis’s cutoff. The transcripts with a value higher than 0.25 are downregulated in RIF-treated samples. In contrast, the transcripts with values lower than -0.25 are upregulated in RIF-treated samples compared to the untreated controls.

**Fig S3.**
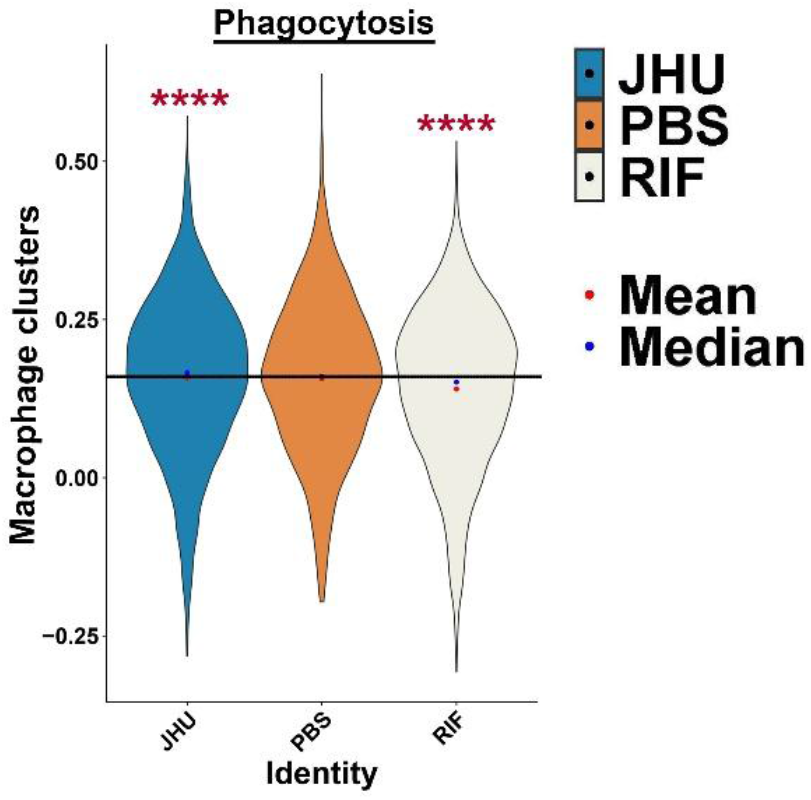
JHU083 administration enriches phagocytosis signature upon macrophage clusters. Violin plot depicting phagocytosis expression score upon macrophage clusters comprising proinflammatory and anti-inflammatory interstitial macrophages. The expression score was calculated using the AddModuleScore module in the R package. The transcripts used for the expression score calculations are listed in **supplementary table 2**. The solid line represents the value corresponding to the mean value of the untreated control samples (PBS). Mean (red dot) and median (blue) values were plotted using ggplot2. ANOVA was used to calculate statistical significance. **** P<0.00001.

**Fig S4.**
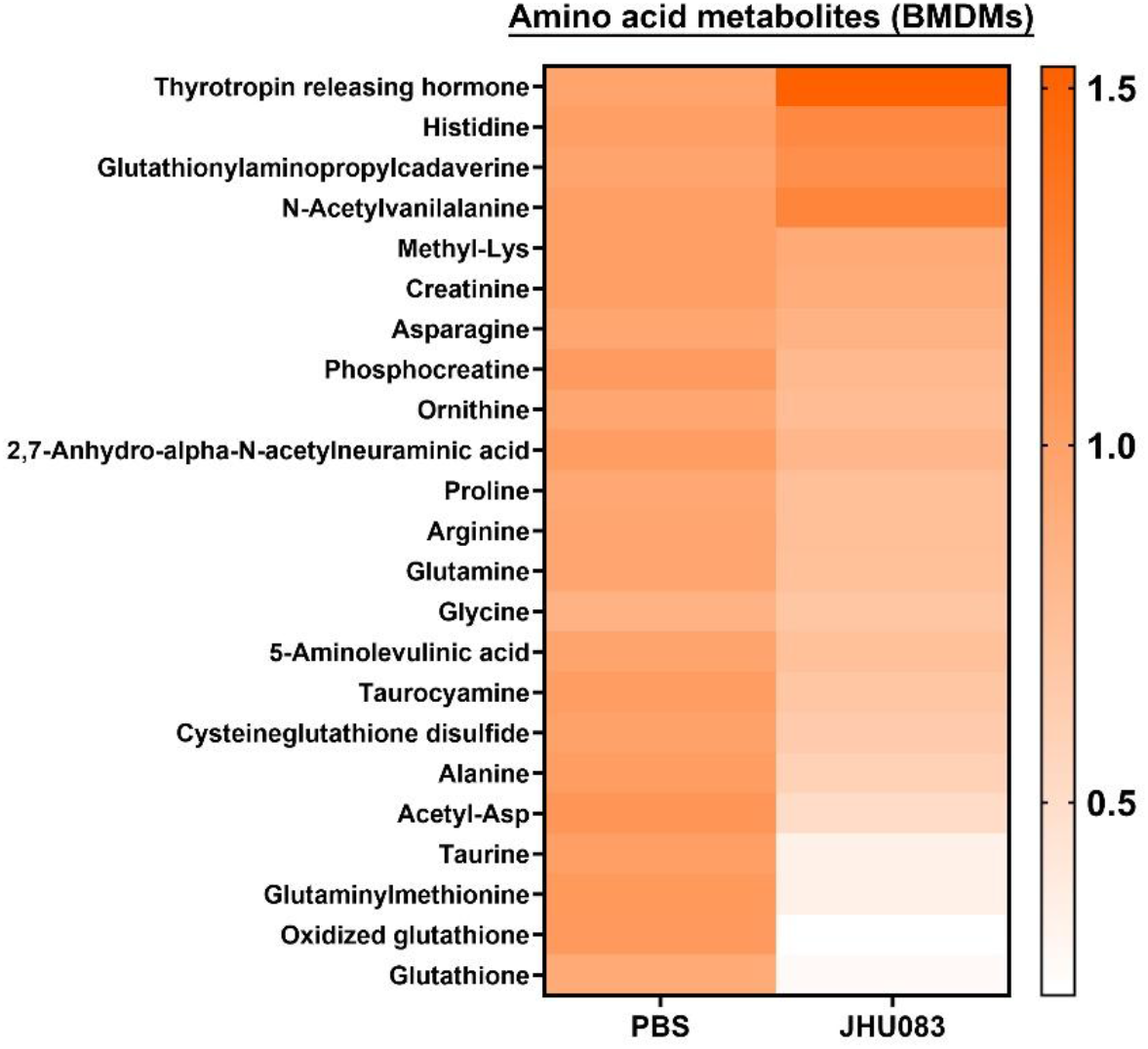
JHU083 treatment depletes amino acids and related metabolites in macrophages. *Mtb*-infected BMDMs were treated with JHU083 or PBS for 24h, and metabolites were extracted from macrophages using methanol as described in the “Methods.” Metabolite levels were normalized to cell number and then to untreated controls. Heatmap representing the top amino acid and amino acid-derived metabolites that were either enriched or depleted in the macrophages treated with JHU083 (right-lane) compared to untreated group (left lane). Metabolite abundance was normalized to the number of macrophages used for extraction and to untreated controls. Statistical significance was determined using a two-tailed Student’s t-test assuming unequal variance. Data are representative of two independent experiments.

