## Supplementary material for "Single-cell RNA sequencing reveals that host Glutamine Metabolism Inhibition Enhances Macrophage Phagocytosis of *Mycobacterium tuberculosis*": Table 1 and 2

**Table 1: The average number of the cells per group**

| | PBS | | | JHU083 | | | RIF | | |
|---|---|---|---|---|---|---|---|---|---|
| **Replicates** | **M1** | **M2** | **M3** | **M1** | **M2** | **M3** | **M1** | **M2** | **M3** |
| **Total cells loaded** | 10,000 | 10,000 | 10,000 | 10,000 | 10,000 | 10,000 | 10,000 | 10,000 | 10,000 |
| **Total cells captured** | 8110 | 7150 | 9396 | 5049 | 10258 | 7107 | 7572 | 5882 | 7881 |
| **Total immune cells** | 7930 | 7015 | 9294 | 4765 | 9656 | 6635 | 7006 | 5126 | 6331 |
| **% Immune cells** | 97.78 | 98.11 | 98.91 | 94.38 | 94.13 | 93.36 | 92.53 | 87.15 | 80.33 |
| **Average % Purity** | | **98.27 ± 0.47** | | | **93.96 ± 0.43** | | | **86.67 ± 4.9** | |

**Table 2: Total 15 clusters identified in the ScRNA seq analysis**

| No | Cluster | Cell types | Markers | Acronym |
|---|---|---|---|---|
| 1 | 1 | Naïve CD4+ T-cells | *Cd3d, Cd4, Lef1, Sell, Ccr7, Slpr1* | CD4_Na |
| 2 | 7 | Naïve CD8+ T-cells | *Cd3d, Cd8a, Lef1, Sell, Ccr7, Slpr1* | CD8_Na |
| 3 | 3 | Exhausted effector CD4+ T-cells | *Cd3d, Cd4, Icos, Rora, Stat4, Stat1, Foxp1, Ctla4* | CD4_Ex |
| 4 | 16 | Terminally exhausted CD4+ T-cells | *Cd3d, Cd4, Klrg1, Cx3cr1, Zeb2, Ifng, Il18r1* | CD4_TEx |
| 5 | 2 | Exhausted CD8+ T-cells | *Cd3d, Cd8aIcos, Rora, Stat4, Stat1, Foxp1, Ctla4* | CD8_Ex |
| 6 | 21 | Terminally Exhausted CD8+ T-cells | *Cd3d, Cd8a, Klrg1, Cx3cr1, Zeb2, Gzma, Il18r1* | CD8_TEx |
| 7 | 9 | Regulatory T cells | *Cd3d, Cd4, Foxp3, Areg, Ctla4, Icos, Il10* | Tregs |
| 8 | 12 | Double-positive T-cells | *Cd3d, Cd4, Cd8a, Cd8b1* | DPT |
| 9 | 0 | B-cells | *Cd19, Cd20, Cd79a, Cd79b, Ms4a1* | B |
| 10 | 8 | Pro-inflammatory M1 macrophages | *Cd14, Csf1, Adgre4, Fcgr4, Dusp16, Lyz2, Pglyrp1* | M1_Mac |
| 11 | 15 | Anti-inflammatory M2 macrophages | *Cd14, Csf1, Adgre4, Chil3, Ccr2, Ccl2* | M2_Mac |
| 12 | 5 | Classical Monocytes | *Cd14, Csf1r, Ifitm3, Apoe, Ly6a, Aif1, Nos2, Cxcl16* | cMo |
| 13 | 11 | Alveolar macrophages | *Siglecf, Marco, Lpl, Mrc1* | AM |
| 14 | 4 | Granulocytic MDSCs (gMDSCs) | *Cd14, Csf3r, Retnlg, S100a8, S100a9, Arg2, Cxcr2* | gMDSC |
| 15 | 14 | Monocytic MDSCs (mMDSCs) | *Cd14, Csf1r, Arg2, Tnf, Ccl3, Ccl4, Il1rn* | mMDSC |
